# A paternal signal induces endosperm proliferation upon fertilization in Arabidopsis

**DOI:** 10.1101/2023.01.12.523779

**Authors:** Sara Simonini, Stefano Bencivenga, Ueli Grossniklaus

**Author notes:** Correspondence: Ueli Grossniklaus, Sara Simonini.

## Abstract

In multicellular organisms, sexual reproduction relies on the formation of highly specialized, differentiated cells, the gametes. At maturity, male and female gametes are quiescent, awaiting fertilization, with their cell cycle being arrested at a precise stage. Failure to establish quiescence leads to unwanted proliferation, abortion of the offspring, and a waste of resources. Upon fertilization, the cell cycle resumes, allowing the newly formed zygote to divide rapidly. Successful development requires that male and female gametes are in the same phase of the cell cycle. The molecular mechanisms that enforce quiescence and reinstate cell division only after fertilization occurs are poorly understood. Here, we describe a sperm-derived signal that induces proliferation of the *Arabidopsis* central cell precisely upon fertilization. We show that the mature central cell is arrested in S phase, caused by the activity of the conserved RETINOBLASTOMA RELATED1 (RBR1) protein. Paternal delivery of the core cell cycle component CYCD7;1 triggers RBR1 degradation, thereby stimulating S phase progression. Absence of CYCD7;1 delays RBR1 depletion, S phase reactivation, and central cell division, whereas its constitutive expression triggers proliferation of unfertilized central cells. In summary, we show that CYCD7;1 is a paternal signal that informs the central cell that fertilization occurred, thus unlocking quiescence and ensuring that cell division initiates just at the right time to ensure functional endosperm formation.

## Introduction

Sexual reproduction entails the specification of the germline and the formation of male and female gametes that fuse during fertilization^1^. In flowering plants (angiosperms), fertilization is unique as it involves two pairs of gametes: the pollen tube delivers two sperm cells (SP) to the female gametophyte, where one fuses with the egg (EC) and the other with the central cell (CC) in a process called double fertilization^2^. Fertilization of the EC and CC result in the zygote and endosperm, respectively, the latter being a triploid placenta-like tissue sustaining embryonic growth^3^. One of the most evident consequences of gamete fusion is the initiation of mitotic divisions upon fertilization. However, in case of inequity between the parental genomes, cell division arrests soon after fertilization, leading to the abortion of the progeny^4–9^. To ensure normal development, male and female gametes establish a quiescent state, such that they do not divide in the absence of fertilization and their cell cycles are synchronized when nuclear fusion (karyogamy) occurs. For instance, mammalian SP and EC are arrested in G1 and in M phase of meiosis II, respectively^10^. Upon fertilization, meiosis is completed such that maternal and paternal genomes are both in G1 phase when embryogenesis starts. In plants, depending on the species, SCs are arrested in G1^11–17^ or G2^18,19^, whereas the cell cycle stage at which the female gametes attain quiescence is still under debate^17,20^.

More than a hundred factors govern cell cycle progression and arrest^21,22^, with their deregulation often leading to reduced fertility because of aberrant gametogenesis and/or embryogenesis^23–27^. Interestingly, in the model plant *Arabidopsis*, mutations in some essential cell cycle genes affect only one of the gametes, with some striking examples affecting the CC where opposite phenotypes, such as proliferation in absence of fertilization or lack of division upon karyogamy, have been observed^28–32^. These findings have fuelled the hypothesis that a mechanism preventing cell division operates in the CC, which is lifted by a fertilization-dependent signal^20,33,34^. Albeit attractive, the molecular players underlying this proposed mechanism are yet to be identified.

## Results

To investigate cell cycle dynamics around fertilization, we first assessed the stage at which mature female gametes arrest. Analysis of the expression levels of components of the cell cycle machinery using available transcriptome datasets^35–37^ for the EC, CC, and pollen (PO) indicated that the CC expresses high levels of virtually all factors involved in and necessary for DNA replication during S phase, while this is not the case for the EC and PO (Fig. 1a). To explore this aspect at the cellular level, we quantified the DNA content in the EC and CC using propidium iodide^38^, which revealed that the unfertilized (virgin) EC and CC have a ploidy level of 1n/2c and about 2n/3c, respectively (Fig. 1b-f), with n indicating chromosome copies and c the number of sister chromatids. For example, 1n/2c indicates a cell with haploid number of chromosomes (1n), where the DNA has been replicated but the two sister chromatids are not yet separated (2c). From our analysis, we conclude that the EC reaches the G2 phase of the cell cycle right before fertilization, whereas the CC is either progressing through or arrested in S phase. To distinguish between these two possibilities, we incubated inflorescences with the nucleotide analogue 5-Ethynyl-2’-deoxyUridine (EdU), which allows the visualization of DNA synthesis *in situ*. The virgin CC did not incorporate EdU even under prolonged incubation time (Fig. 1g,i), indicating the absence of DNA synthesis. We confirmed this observation by monitoring the expression of the replication licensing factor CTD1a-GFP, whose degradation is promoted by entry and progression through S phase^39,40^. CTD1a-GFP accumulated in the virgin CC (Fig. 1j-l,n), being detectable already in the two polar nuclei in the female gametophyte before they fuse (Fig. 1j-k). Altogether, these data suggest that the virgin CC is arrested in S phase at maturity.

**Figure 1.**
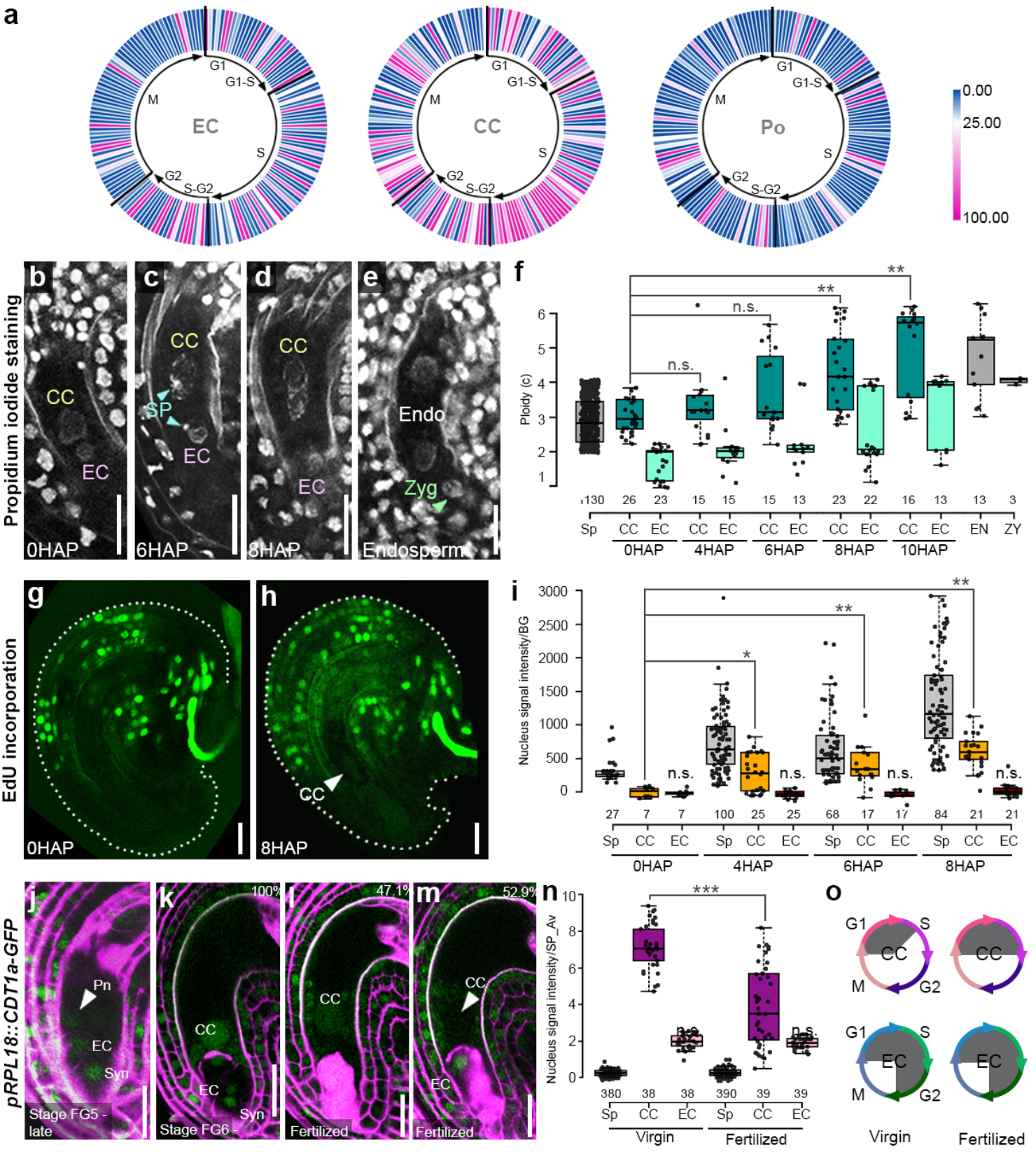
Cell cycle dynamic around the moment of fertilization in the EC and CC. **a.** Circular heat map of the expression level of cell cycle components of Egg Cell (EC), Central Cell (CC), and Pollen (PO). The genes are organized following the cell cycle stages, from G1 to M. A single gene can be represented in more than one stage, accordingly to its function. **b-e.** Representative images of Propidium Iodide staining of ovules, with a focus on the whole female gametophyte, at different time points after pollination: 0HAP (b), 6HAP (c), 8HAP (d), after the first endosperm division (e). **f.** Box plot showing Propidium Iodide staining quantification as ploidy value of sporophytic cells (Sp), CC, EC, Endosperm (EN), and zygote (ZY), at different time point after pollination. n.s., not significant; * P-value<0.01, ** P-value<0.001 accordingly to a *t*-test. **g-h.** Representative images of EdU staining of ovules at 0HAP (g) and 8HAP (h). **i.** Box plot showing EdU signal quantification in sporophytic cells (Sp), CC, and EC, at different time points after pollination. n.s., not significant; * P-value<0.01, ** P-value<0.001 accordingly to a *t*-test. **j-m.** Confocal analyses of CTD1a-GFP dynamic in the female gametophyte at stage late FG5 (j), stage FG6 (k), and in fertilized ovules (l-m). Cell walls are stained with Propidium Iodide. Fertilized ovules are easily identified as they accumulate Propidium Iodide in the penetrated synergid cell. **n.** Quantification of the signal intensity of CTD1a-GFP in Sp, EC, CCs in virgin and fertilized ovules. Values for each category are normalized on the averaged signal intensity value of ten sporophytic cells surrounding the female gametophyte. n.s., not significant; *** P-value<0.0001 according to a *t*-test. **o.** Schematic representation summarizing the cell cycle dynamic of EC and CC according to the fertilization event. Virgin and mature ECs are in G2 phase, virgin CCs are in arrested S phase, whereas fertilized CCs exhibit S phase reactivation. Scale bar: 20μm.

After fertilization, we observed remarkable changes in the CC. Firstly, the DNA level increased from 2n/3c to about 3n/5c, indicating successful fertilization (Fig. 1f and Extended Data Fig. 1a) and supporting previous observations that the sperm cell has a DNA amount of 1n/2c^18^. At about eight <u>h</u>ours <u>a</u>fter <u>p</u>ollination (HAP), the fertilized CC reached a ploidy level of 3n/6c, indicating that cell was now in G2 phase (Fig. 1f and Extended Data Fig. 1a). Nuclear fusion between SPs and EC or CC happens independently of each other^34,41–44^. Indeed, we detected ovules where the EC showed a ploidy level of 2n/4c, while the CC still exhibited one of 2n/3c (Extended Data Fig. 1a), with a visible SP nucleus in the proximity. This observation indicates that SP delivery to the female gametophyte is not enough to trigger cell cycle reactivation in the CC but, instead, that karyogamy is required, as an increase of the ploidy level in the CC occurs only after it reached a value of 3n/5c. In addition to the increase of the ploidy level occurring after karyogamy, we also observed faint but detectable EdU staining in the CC (Fig. 1h-i), and depletion of CTD1a-GFP exclusively form the CC (Fig. 1m-o), with both events occurring only in fertilized ovules. The EC behaved very differently: following fertilization, its ploidy level sharply increased from 1n/2c to 2n/4c (Fig. 1f and Extended Data Fig. 1a), without detectable EdU incorporation (Fig. 1i) and persisting CTD1a-GFP signal (Fig. 1n). Altogether, these observations confirm that the mature EC and SP are arrested at the G2 phase of the cell cycle^18^, and revealed that the CC is arrested during S phase, which is completed only after karyogamy (Fig. 1o).

Next, we sought to investigate the mechanism underlying karyogamy-dependent S phase reactivation in the CC by focusing on the core cell cycle regulator RETINOBLASTOMA RELATED1 (RBR1)^45–48^. RBR1 is a potent inhibitor of entry into and progression through S phase^49^, and *rbr1* mutant CCs exhibit uncontrolled divisions to produce an endosperm-like structure^29^ (Fig. 2a-b), suggesting that RBR1 may be required for S phase arrest in the CC. Therefore, we investigated RBR1 protein dynamics in the CC around fertilization. Live imaging analysis of *pRBR1::RBR1-YFP*^50^ in unfertilized ovules confirmed the previously characterized accumulation of RBR1 in the virgin CC^51^ (Fig. 2c-d). However, around 7-8 HAP, the *pRBR1::RBR1-YFP* signal in the CC became progressively weaker and finally undetectable, before reappearing in the first two endosperm nuclei (first CC division; Fig. 2c-g). To investigate whether RBR1 degradation in the CC is mediated the 26S proteasome, as it occurs in animals and in other plant tissues^50,52^, we treated pistils with the selective 26S proteasome inhibitors Epoxomycin, Syringolin-A (SylA), and MG-132, and repeated the live imaging analysis of *pRBR1::RBR1-YFP* around fertilization (Fig. 2h). Upon Epoxomycin and SylA treatment the *pRBR1::RBR1-YFP* signal persisted in the fertilized CC (Fig. 2h) in comparison to MOCK-treated pistils (Fig. 2h), thereby preventing - or slowing down - *pRBR1::RBR1-YFP* degradation in the CC nucleus. Although MG-132 is a known inhibitor of RBR1 degradation^50^, it did not have significant effects on *pRBR1::RBR1-YFP* stability in the CC under our experimental conditions (Fig. 2h). A possible explanation lies in the chemical properties of the inhibitors: MG-132 is a reversible inhibitor that is highly unstable in aqueous solutions, whereas Epoxomycin and Syl-A are high-affinity irreversible, water-stable inhibitors. We repeated Epoxomycin and Syl-A treatments and left the pollinated pistils to develop for a further four days. Remarkably, a single treatment was sufficient to induce the development of seeds where normal-looking globular stage embryos were surrounded by a few, massively enlarged endosperm nuclei (Fig. 2i-k). Similar endosperm phenotypes also develop in plants lacking subunits of the DNA replication machinery, such as ORIGIN OF REPLICATION COMPLEX2^32^ (ORC2), MICROSOMAL MAINTENANCE2^30^ (MCM2), MCM7/PRL^31^, and CULLIN4 (CUL4)^53^, an E3 ubiquitin-ligase involved in protein degradation through the 26S proteasome. Altogether, these data suggest that in the virgin CC, the persistence of RBR1 leads to an arrest of the cell cycle in S phase. Moreover, removal of RBR1 through the 26S proteasome-mediated degradation pathway is a pre-requisite for S phase completion, ultimately ensuring synchronized maternal and paternal genomes to initiate endosperm development.

**Figure 2.**
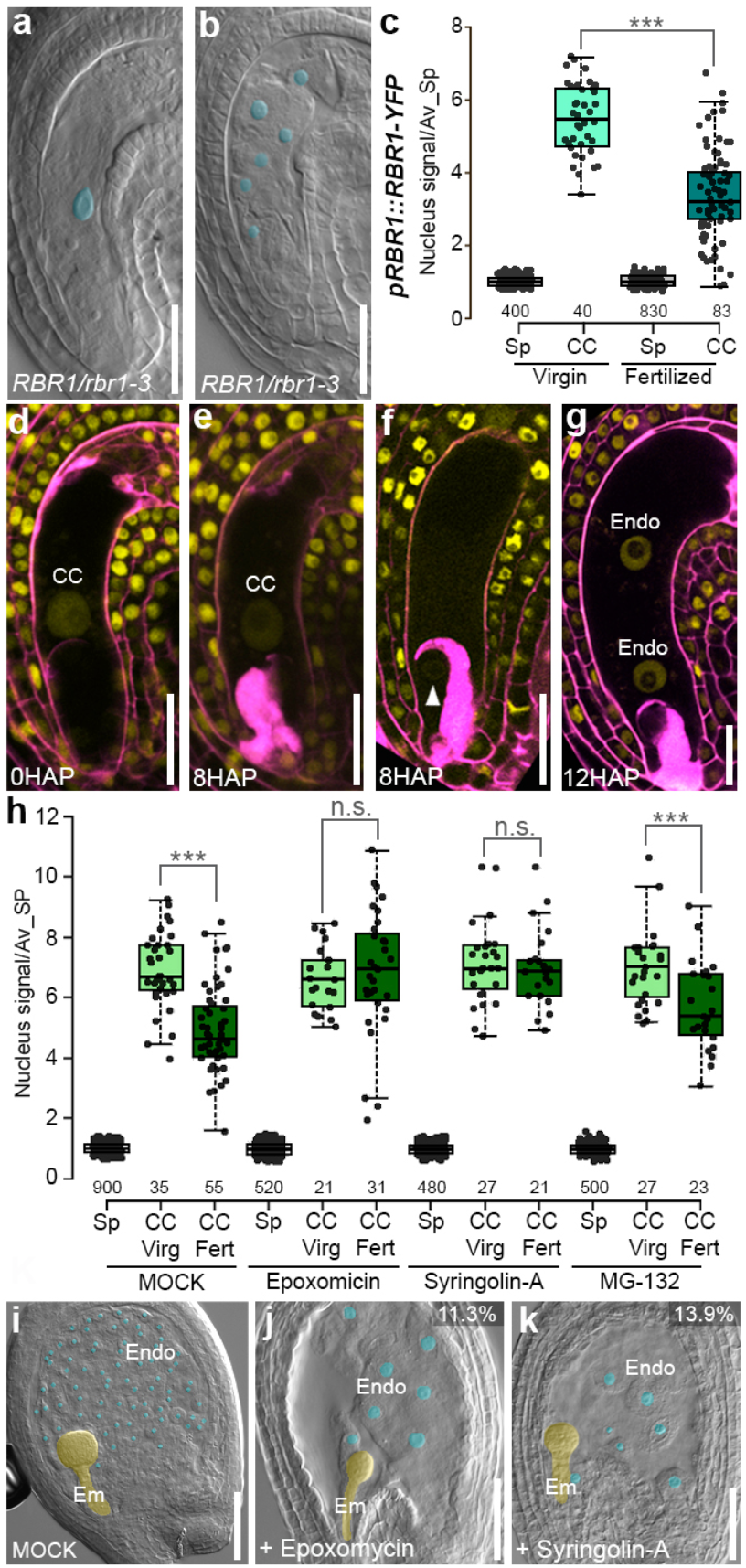
RBR1 protein is depleted from the CC nucleus after fertilization through 26S-proteasomal degradation. **a-b.** Clearing analysis of virgin ovules from *RBR1/rbr1-3* plants, showing a wild-type female gametophyte (a), and a mutant female gametophyte (b) where the CC has undergone division. The CC nucleolus in (a) and the endosperm-like nucleoli in (b) have been artificially highlighted in cyan. **c.** Quantification of the RBR1-YFP signal intensity in sporophytic cells (Sp) and CCs of virgin and fertilized ovules. Values for each category are normalized on the averaged signal intensity value of ten Sp surrounding the female gametophyte. *** P-value<0.0001 according to a *t-* test. **d-g**. Representative images of *pRBR1::RBR1-YFP* ovules at different time points after pollination: 0 HAP (d), 8 HAP (e-f), and 12 HAP (g). Image in (f) has increased contrast to reveal RBR1-YFP signal in the synergid nucleus (white arrowhead). **h.** Quantification *of pRBR1::RBR1-YFP* signal in Sp, virgin and fertilized CCs, from inflorescence treated with a MOCK solution, and the proteasome inhibitors Epoxomycin, Syl-A, and MG-132. Values for each category are normalized on the averaged signal intensity value of ten Sp surrounding the female gametophyte. n.s., not significant; *** P-value<0.0001 according to a *t*-test. **i-k.** Clearing analysis of seeds from inflorescences treated with a MOCK solution (i), Epoxomycin (j), and Syl-A (k). At the top right corner is the percentage of seeds showing the phenotype. Embryo and endosperm nucleoli are artificially highlighted in yellow and cyan, respectively. Scale bar: 20μm a-b, c-g; 50μm i-k.

To understand how RBR1 degradation is specifically achieved at karyogamy in the CC, we performed transcriptome analyses of CCs isolated by Laser-Assisted Microdissection (LAM) at different time points (Fig. 3a, Extended Data Table 1 and Extended Data Fig. 2a-d). Stages encompassed 0 HAP (virgin CC), 4 HAP (fertilization occurred in about 50% of the CCs), 8 HAP (all CCs fertilized), and 12 HAP (around the first endosperm division). Our goal was to identify transcripts whose expression peaked around 8 HAP, the time point at which RBR1 degradation occurs in the CC, to differentiate between two possible scenarios: either their transcription occurred *de novo* in a fertilization-dependent manner, or they were of paternal origin and delivered to the CC upon gamete fusion. Since phosphorylation-dependent RBR1 degradation occurs by a cell cycle-regulated mechanism^49,54–56^, we focused on cell cycle components whose expression changed across the four time points (Fig. 3b). One candidate belonging to the Cyclin D-type family, *CYCD7;1*, captured our attention for several reasons: (i) D-type cyclins mediate entry and progression through S phase^57,58^; (ii) CYCD7;1 interacts with RBR1 and promotes its degradation^59,60^; (iii) *CYCD7;1* transcript was absent in virgin CCs, sharply increased at 8 HAP (Fig. 3b), and was the only D-type cyclin with high expression in pollen^37,59^, and (iv) ectopic expression of *CYCD7;1* in ovules can induce CC proliferation in absence of fertilization^61^. Based on these observations, we hypothesize that CYCD7;1 serves as a paternal signal that is loaded into the SP and delivered to the CC upon fertilization, thus provoking RBR1 degradation and triggering S phase completion. To confirm this attractive hypothesis, we first investigated the origin of the *CYCD7;1* mRNA by *in-situ* hybridization of wild-type pistils fertilized with pollen from a *pCYCD7:1:;CYCD7;1-YFP* line^59^, using an antisense probe specific to *YFP* (Fig. 3c-j). A signal was observed in the *pCYCD7:l:;CYCD7;1-YFP* line during pollen development (Fig. 3d) and in the SP nucleus of both mature pollen grains and in elongating pollen tubes (Fig. 3e-f). A strong, punctate signal was also observed in wild-type ovules fertilized with *pCYCD7:1:;CYCD7;1-YFP* pollen at about 4 HAP, labelling the discharged SP nucleus (Fig. 3h). Afterwards, the signal was detected in the CC, initially with a precise nuclear confinement (4 HAP, Fig. 3i), and later also in the CC (and EC) cytoplasm (Fig. 3j). To further corroborate that *de novo CYCD7;1* transcription does not occur in the CC, neither from the maternal nor the paternal allele, we analysed the expression pattern of the transcriptional reporter *pCYCD7;1::YFP-YFPnls*^59^. The YFP signal was detected sporadically in the synergid cells of unfertilized ovules (Fig. 3k) but not in the virgin or fertilized CCs and ECs (Fig. 3l).

**Figure 3.**
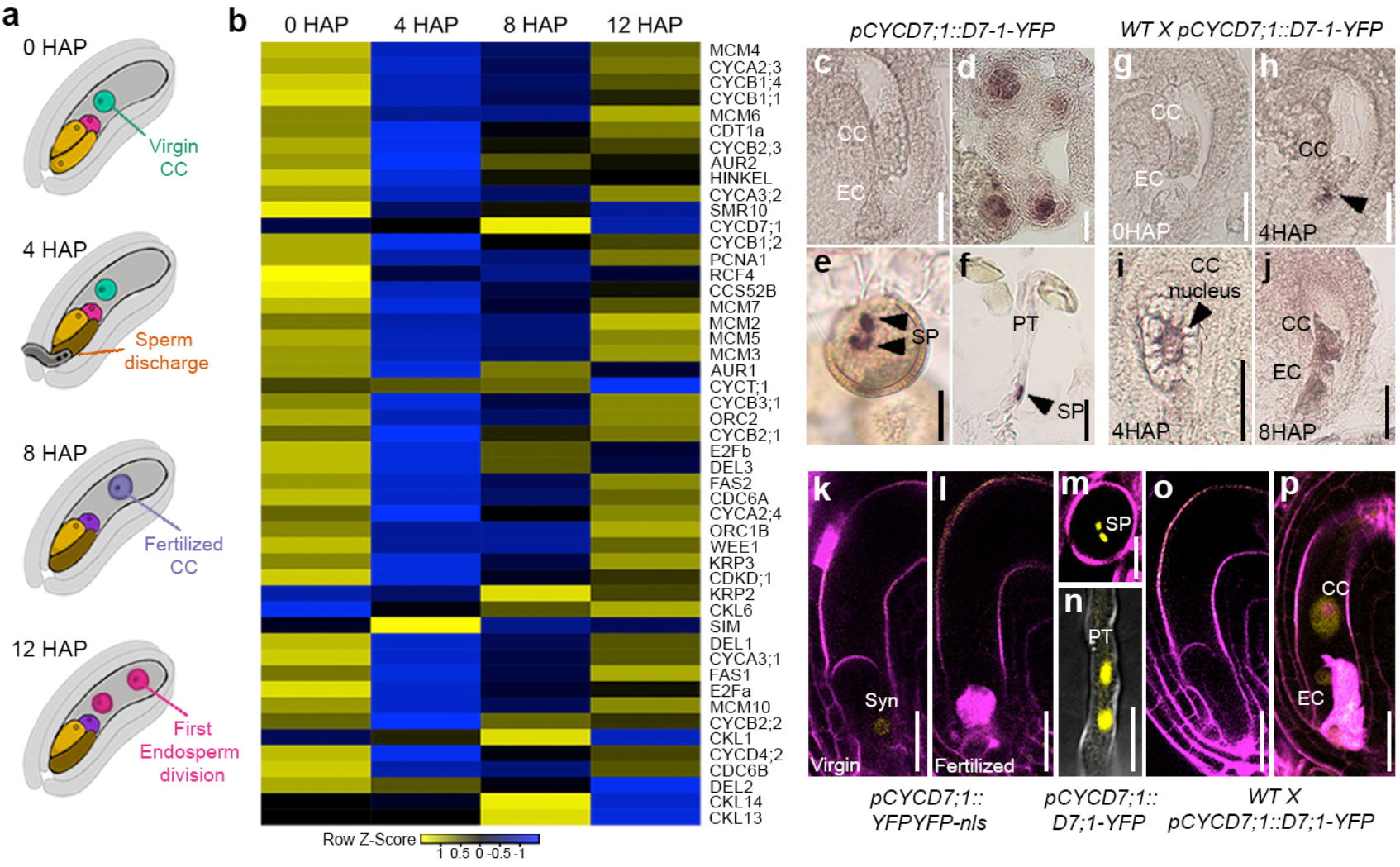
*CYCD7;1* mRNA and protein have paternal origin and are delivered to the CC at fertilization through karyogamy. **a.** Graphic representation of the strategy adopted for the LCM transcriptome analysis of CC at different time points: 0 HAP, 4 HAP, 8 HAP, and 12 HAP. **b.** Heat map with cell cycle-related genes that show variation of expression across the four LCM data points. **c-f.** *In-situ* hybridization with a YFP antisense probe on *pCYCD7;1::CYCD7;1-YFP* plants showing no signal in virgin ovules (c), but a positive signal in anthers (d), the SP nuclei in mature pollen grains (e) and within growing pollen tubes (f). **g-j.** *In-situ* hybridization with a YFP antisense probe on wild-type pistils pollinated with *pCYCD7:1::CYCD7;1-YFP* pollen at 0 HAP (g), 4 HAP (h-i) and 8 HAP (j), with a signal in the discharged SPs (arrowhead, g), the CC nucleus (i), and CC and EC nuclei and cytosols (j). **k-l.** Confocal images of *pCYCD7;1::YFPYFP-nls* in virgin (k) and fertilized (l) ovules. **m-n.** Confocal analysis of the *pCYCD7;1::CYCD7;1-YFP* line in mature pollen grains (m), and pollen tubes (n) showing positive signal in the SP nuclei. **o-p.** Confocal analysis of virgin (o) and pollinated (p) wild-type ovules with *pCYCD7;1::CYCD7;1-YFP* pollen showing a positive signal in the fertilized CC and EC. CC, Central Cell; EC, Egg Cell; PT, Pollen Tube; Syn, Synergid cell; SP, Sperm cell. Scale bar: 10μm e,I,n, otherwise 20μm.

Previous work demonstrated that CYCD7;1 protein expression is restricted to the stomatal lineage and mature pollen^59,62^. Therefore, we wondered whether, in addition to its mRNA, also the CYCD7;1 protein may be stored in the SP nucleus and delivered to the female gametophyte. Analysis of *pCYCD7:1:;CYCD7;1-YFP* confirmed this hypothesis: CYCD7;1-YFP-expressing SP nuclei were visible in mature pollen grains (Fig. 3m), SP nuclei inside growing pollen tubes (Fig. 3n), and the fertilized CC and EC nuclei (Fig. 3o-p). Altogether, these results demonstrate that CYCD7;1 is a paternally derived factor, with its transcripts and protein being delivered directly to the nuclei of the female gametes. We propose that this specific, restricted nuclear localization of mRNA and protein secures that CYCD7;1 is active only upon successful karyogamy, and not simply after SP delivery.

Next, we questioned whether CYCD7;1 promotes RBR1 degradation in the CC. Pollination of *pRBR1::mCherry-RBR1^63^* pistils with *pCYCD7;1::CYCD7;1-YFP* pollen showed that RBR1 depletion in the CC indeed occurs after CYCD7;1 delivery (Extended Data Fig. 3a-c). Moreover, ectopic expression of *CYCD7;1* in the CC under a CC-specific^64^ *(pMEA::CYCD7;1;* n=28/30 independent transformants, Fig. 4a-b) or ubiquitously expressed promoter *(pRPL18::CYCD7;1*, n=18/26 independent transformants, Fig. 4c), respectively, induced the development of endosperm-like structure in virgin ovules, as previously reported^61^. The nuclei showed a very weak or undetectable *pRBR1::RBR1-YFP* signal (Fig. 4e-f), indicating that CYCD7;1 alone is sufficient to induce RBR1 degradation in the CC and, consequently, to stimulate S phase progression. Because the interaction between RBR1 and CYCD7;1 is mediated by the Leu-x-Cys-x-Glu (LxCxE) motif^60,65^, we ectopically expressed a CYCD7;1 LxCxE mutant variant *(pRPL18::CYCD7;1^mut^)*. Contrarily to the wild-type version, CYCD7;1^mut^ was incapable of inducing CC proliferation in virgin ovules (n=58; 0% of endosperm-like structures detected, Fig. 4d), suggesting that the interaction of CYCD7;1 with RBR1 is required to mediate S phase progression in the CC upon fertilization.

**Figure 4.**
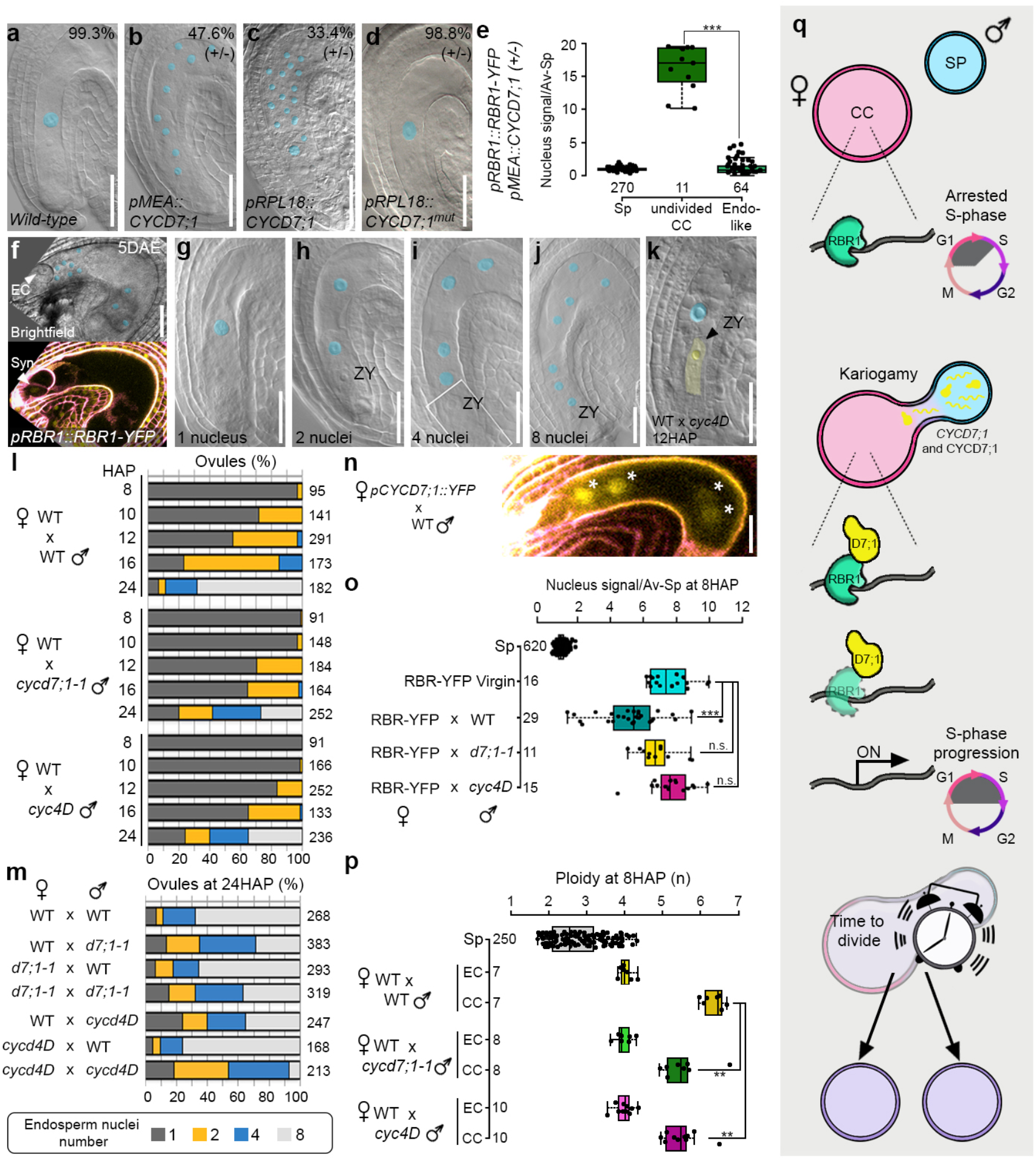
CYCD7;1 is paternally required for timely RBR1 degradation and cell division in the fertilized CC. **a-b.** Clearing analysis of ovules from plants heterozygous for the *pMEA::CYCD7;1* construct, displaying a wild-type looking ovule with an undivided CC (a), and an ovule with CC proliferation. In the top right corner is the percentage of ovules showing the corresponding phenotype. **c-d.** Clearing analysis of ovules of heterozygous plants expressing the *pRPL18::CYCD7;1* and *pRPL18::CYCD7;1^mut^* constructs, showing an ovule whit CC proliferation (c) and wild-type looking ovules (d). In the top right corner is the percentage of ovules showing the corresponding phenotype. **e.** Quantification of the *pRBR1::RBR1-YFP* signal in the *pMEA::CYCD7;1/-* background at 5 days post-emasculation, in undivided CC and endosperm-like nuclei. For each ovule showing CC proliferation, 4 nuclei have been selected *** P-value <0.0001 accordingly to a *t*-test. **f.** Example image of the *pRBR1::RBR1-YFP* signal in the *pMEA::CYCD7;1/-* background in a ovule that showed CC proliferation in absence of fertilization. **g-j**. Clearing analysis of fertilized ovules showing four endosperm stages: fertilized CC (d), two (e), four (f) and eight (g) endosperm nuclei. **k.** Clearing analysis of a wild-type ovule fertilized with *cycd4D* pollen at 12 HAP, showing an undivided CC in proximity of the elongating zygote. **l.** Classification of fertilized ovules (in percentage) based on the number of endosperm nuclei at a certain time point (8, 10, 12, 16, 24 HAP), in crosses between wild-type (*WT) x WT, WT x cycd7;1-1*, and *WT x cyc4D*. Sample size is indicated on the right side of each bar. The colour code is at the bottom of the panel. **m.** Classification of fertilized ovules (in percentage) based on the number of endosperm nuclei at 24 HAP, in reciprocal crosses between *WT, cycd7;1-1*, and *cyc4D* individuals. Sample size is indicated on the right side of each bar. **n.** Confocal analysis of the *pCYCD7;1::YFPYFPnls* line at 16HAP in reciprocal crosses with WT plants showing activation of the maternal *pCYCD7;1::YFPYFPnls* transgene. **o.** Box plot showing the quantification of *pRBR1::RBR1-YFP* signal in virgin CC, and in RBR1-YFP pistils pollinated with WT, *cycd7;1* or *cyc4D* pollen. n.s., not significant; *** P-value<0.0001 accordingly to a *t*-test. **p**. Quantification of ploidy value through Propidium Iodide staining of CC and EC at 8 HAP of WT pistils pollinated with WT, *cycd7;1* or *cyc4D* pollen. ** P-value<0.001 accordingly to a *t*-test. **o.** Graphic representation summarizing the final model proposed in this work: delivery of paternally originated CYCD7;1 mRNA and protein at fertilization triggers completion of the S phase in the CC through RBR1 depletion, thereby allowing S phase progression to allow timely cell division in the endosperm. Scale bar: 20μm

To investigate whether CYCD7;1 exerts a paternal control over cell division in the fertilized CC, we characterized *CYCD7;1* T-DNA insertion mutants *(cycd7;1-1^59^, cycd7;1-2*^59^, and *cycd7;1-3)* and new mutant alleles created by CRISPR-Cas9 technology *(cycd7;1^CRISPR^*, Extended Data Fig. 3d). We focused on CC division by scoring the percentage of fertilized ovules with an undivided CC (Fig. 4g), or two, four, or eight endosperm nuclei (Fig. 4h-j). In reciprocal crosses between wild-type pistils and *cycd7;1* mutant pollen, we observed a significant delay of the first CC division (Fig. 4l and Extended Data Fig. 3e). At 10 HAP, wild-type x wild-type crosses showed two endosperm nuclei in about 30% of the fertilized ovules (Fig. 4h,l and Extended Data Fig. 3h), while in seeds originating from crosses between wild-type x *cycd7;1* mutants, less than 5% had two endosperm nuclei (Fig. 4l and Extended Data Fig. 3e). We confirmed that the delay in endosperm initiation was not caused by delayed fertilization, as wild-type and *cycd7;1* mutant pollen tubes had comparable growth rates (Extended Data Fig. 4a). Because seeds receiving a paternal *cycd7;1* mutant allele started to recover around 16 HAP (Fig. 4i), we investigated whether other D-type cyclins could compensate for the absence of *CYCD7;1* and, therefore, created the quadruple mutant *cycd3;1 cycd3;2 cycd3;3 cycd7;1* (referred to as *cyc4D;* Extended Data Fig. 4b-c). Wild-type pistils pollinated with *cyc4D* pollen showed an even longer delay of the first endosperm division (99% of ovules at 10 HAP still had an undivided CC; Fig. 4h-i). Embryo development was not affected, and we detected several fertilized ovules with an undivided CC nucleus next to an elongating zygote (Fig. 4k). Remarkably, the delay in initiation of the first endosperm division was further exacerbated when *cyc4D* pistils were pollinated with *cyc4D* pollen (Figure 4m), suggesting that the fertilized CC becomes transcriptionally active within few hours after fertilization producing cell cycle components.

In support of this hypothesis, we detected a *pCYCD7:l::YFP-YFPnls* signal from the maternal allele in 4-nuclear endosperm (around 20 HAP, Fig. 4n), indicating activation of *CYCD7;1* transcription. *CYCD7;1* is also among the cell cycle components that show transcriptional activation from the maternal genome soon after fertilization in the zygote^66^.

Finally, we investigated RBR1 protein dynamics and the ploidy level of the fertilized CC when wild-type, *cycd7;1*, or *cyc4D* plants were used as pollen donors. We detected depletion of *pRBR1::RBR1-YFP* in the fertilized CC at around 8 HAP only when using wild-type pollen, while ovules that received *cycd7;1* or *cycd4D* mutant SPs showed persistence of the RBR1-YFP signal (Fig. 4o). Consistent with this finding, the stability of RBR1-YFP protein is accompanied by a lack of S phase progression: CCs of ovules that received *cycd7;1* or *cyc4D* SPs failed to precisely reactivate S phase after fertilization. In these cases, the CC ploidy remained at a level of about 3n/5c at 8 HAP, a time point when all CCs that were fertilized by wild-type pollen had reached the 3n/6c value (Fig. 4p). Therefore, in absence of the paternal delivery of CYCD7;1, RBR1 depletion fails to occur in the CC at the time of fertilization, thus delaying S phase reactivation and progress.

## DISCUSSION

We describe a simple, yet effective, molecular mechanism that relies on a paternal signal to ensure that cell cycle reactivation occurs precisely upon fertilization. Given its fundamental importance for seed development, a tight control of endosperm proliferation is essential to maximise successful reproduction. Interestingly, cell cycle arrest in S phase and its release by the RBR1-CYCD7;1 module seems to occur only in the CC. Indeed, the EC arrests in the G2 phase and, although it expresses *RBR1*, does not rely on CYCD7;1 delivery by the SC to initiate the first mitotic division of the zygote. This suggests that CC and EC, although genetically identical, adopt different pathways to integrate cell cycle progression with developmental programs that may rely on precisely tailored expression levels and flavours of the various cell cycle components. For instance, different from the EC, the CC expresses all players necessary to promote cell division, such as factors involved in DNA replication and the G2-M transition, but also inhibitors of cell cycle progression, for instance members of the KIP-RELATED PROTEIN (KRP) family that prevent G1-S progression (Fig. 1a). Thus, given the expression of many - even antagonistic - cell cycle components, the regulation of cell division in the CC likely depends on the regulation of protein activity rather than its expression as we showed to be the case for RBR1.

To synchronize its phase of the cell cycle with that of the SC, which is arrested in G2 phase, the CC relies on the delivery of a SC-derived signal that marks successful fertilization. RBR1 can only be degraded by the 26S proteasome once it gets phosphorylated by a cyclin-dependent kinase (CDK), thereby releasing the break on S phase entry and progression. It is tempting to speculate that, upon delivery of CYCD7;1 by the SC, an active CDK-CYCD7;1 complex is formed that initiates RBR1 degradation. Once S phase is completed, the cell cycle stage of the CC matches that of the SC and nuclear endosperm proliferation ensues.

To date, only two factors essential to normal seed development are known to show a specific requirement in one of the gametes. The transcripts of *SHORT SUSPENSOR* accumulate in the SC, are delivered to the EC and CC, and control the development of the embryonic suspensor^67^. Peptides of the EMBRYO SURROUNDING FACTOR1 family, on the other hand, accumulate in the CC and non-autonomously control suspensor development^68^. CYCD7;1 is the first described paternal, SC-derived signal that specifically controls CC proliferation and thus endosperm formation.

Our findings answer a fundamental question in developmental biology, namely how a cell determines when it is the right time to divide, a question of that is of particular importance to female gametes as embryogenesis or endosperm development fail if division occurs prior to successful fertilization. At the same time, new questions arise. Indeed, it would be interesting to investigate how the *CYCD7;1* messenger is specifically retained in the nucleus, and through what mechanism the CC arrests in the middle of S phase. Given the conserved role of the factors involved, the understanding of the fertilization-dependent molecular mechanisms that controls endosperm initiation could help in the design of strategies to manipulate seed development in crop species.

## SUPPLEMENTAL INFORMATION

Supplemental Information can be found online at…

## ACKNOWLEDGEMENTS

We thank Ben Scheres (Wageningen University and Research), Dominique Bergmann (Stanford University), Crisanto Gutierrez (Centro de Biología Molecular Severo Ochoa), Arp Schnittger (University of Hamburg), the Nottingham *Arabidopsis* Stock Center, and the *Arabidopsis* Stock Center IJPB-INRAE for providing seeds, Mortiz Nowack (VIB-Ghent) and Crisanto Gutierrez for helpful comments, Celia Baroux (University of Zurich) for suggestions on image acquisition, Christof Eichenberger, Frédérique Pasquer, Arturo Bolaños, Daniela Guthörl, and Peter Kopf (University of Zurich) for general lab support, the CHERI-T meeting for helpful discussions, and Marc W. Schmid (MWSchmid GmbH) for analysis, figure preparation, and deposition of the LCM-RNA-Seq data. We thank Zsuzsanna Hasenkamp and Robert Dudler (University of Zurich) for the gift of Syl-A. This work was supported by the University of Zurich, a postdoctoral fellowship from the European Union’s Marie Skłodowska-Curie Action to SB, and a grant from the Swiss National Science Foundation to UG.

## AUTHOR CONTRIBUTIONS

UG conceived, supervised, and raised funding for the project. SS conceived, designed, and performed the experiments. SS and UG analysed and interpreted the data. SB generated the CRISPR-Cas9 mutants, overexpression lines, and provided technical support for the LCM RNA-Seq. SS and UG wrote, and SB commented on the manuscript.

## MATERIALS AND METHODS

### Plant material and growth conditions

Seeds were sown on half-strength MS media (1/2 MS salt base [Carolina Biologicals, USA], 1% sucrose, 0.05%MES, 0.8% Phytoagar [Duchefa], pH>5.7 with KOH), stratified for 3-4 days at 4°C in the dark, and then moved to long-day conditions (8h dark at 18°C, 16h light at 22°C, 70% humidity). When showing four true leaves, seedlings were transplanted to soil and grown under long-day conditions in a walk-in growth chamber (8h dark, 16h light, 22°C, 70% humidity). Marker lines used in this study are: *pRBR1::RBR1-Venus^50^, pCYCD7;1::CYCD7;1-YFP^59,62^, pCYCD7;1::YFP-YFPnls^59^, pRBR1::mCherry-RBR1*^63^. Mutant alleles for *CYCD7;1* were obtained from NASC and INRA/Versailles (Versailles, France) collections: *cycd7;1-1* (FLAG_369E0259), *cycd7;1-2* (FLAG_498H0859), *cycd7;1-3* (GK_496G06). Mutants for *cycd3;1* (GK_529C07) and *cyd3;3* (GK-169B01) were obtained from NASC.

When performing reciprocal crosses, flowers at stage 11 of development were emasculated, and left 24h to allow complete maturation of the ovules. Pollination has been done always around 8am, and samples collected, fixed, or imaged at the desired time points in the afternoon/evening. Exception was for the 16HAP time point, were pollination has been performed around 17.00, to avoid collection of material during the night.

### Vectors

For construction of *pRPL18::CYCD7;1*, the *pRPL18* and the *CYCD7;1* genomic sequence were amplified from Col-0 genomic DNA using respectively the couple of primers 169-170 and 528-525. The PCR fragments were cloned into the miniT vector (NEB), and assembled together in a Golden Gate reaction together with the terminator *tHSP18.2* (Addgene GB0035) and the destination vector p140 using the enzyme BsaI. The destination vector p140 is a modified version of the Golden Gate low copy binary vector pAGM53451^69^. For our purpose, we inserted in the pAGM53451 vector the RedSeed selector marker at position three, whereas at position two we inserted the *lacZ* gene adapted to be an acceptor for Level 1 Golden Gate modules.

For the *pRPL18::CYCD7;1^mut^*, we mutated the LxCxE motif to XxXxX, accordingly to Weimer et al., 2018^59^, using the same strategy as above by amplifying the *CYCD7;1* gene with primer 577-525, where the former contained the mutations.

For the *pMEA::CYCD7;1*, a 4.5Kb fragment of the *MEA* promoter was amplified with primers M1 and M2 and cloned in miniT plasmid in a similar strategy adopted in Simonini et al., 2021^23^. The construct was assembled as describe before using the Golden Gate system together with and the *miniT-CYCD7;1*, the terminator *tHSP18.2*, and the backbone *p140*.

For *pRPL18::CTD1a-GFP*, the *CTD1a* gene was amplified in two parts from Col-0 genomic DNA using the primers 351-345 and 346-352, cloned in the miniT plasmid (NEB), and the assemble with the Golden gate system using the modules for **<u>GFP</u>**, terminator *tHSP18*, and the destination vector *p140*.

### CRISPR-Cas9

To create the CRISPR construct it has been used the following strategy. Specific gRNA have been designed using the software Chopchop^70^ (https://chopchop.cbu.uib.no) These primers have been used as forward primer specific for the gene of interest together with the reverse primer critarrev (atgtacggccagcaacgtcg) using as template the plasmid pAGM9037. Next a golden gate reaction has been done to insert the PCR product in the MoClo level 1 plasmid^71^ at the position 3 (when only one gRNA) or 3 and 4 when two gRNA were required for the construct. The level 1 plasmid with gRNA have been then assemble in a level 2 plasmid pAGM65879 together with the selection in plant for red florescence seed coat. The final construct has been transformed in *A.tumefaciens* GV3101 and then in plants. T1 seeds have been selected for red fluorescence in seed coat. In generation T2 only the seeds without red fluorescence have been picked in order to select plants without the construct.

### Imaging and image analysis

All imaging of *pRBR1::RBR1-YFP* line has been performed using a Leica TCS SP8 Multiphoton microscope, equipped with RLD detectors, and in photon counting mode. Z-stacks that include the entire embryo sac have been acquired. Then, the focal planes encompassing CC, EC or sporophytic cells have been merged in a single plane image using the Sum command in Fiji-ImageJ. For each ovule, 10 sporophytic cells of the inner integument surrounding the embryo sac have been measured to normalize the signal intensity of the CC and EC. Nuclei have been manually identified.

### Proteasome inhibitor treatment

Inflorescences were cut leaving 1.5cm stem, and inserted in a 2ml cryotube filled with MS solid media (1/2 MS salt base [Carolina Biologicals, USA], 1% sucrose, 0.05%MES, 0.8% Phytoagar [Duchefa], pH>5.7 with KOH), supplemented with the desired inhibitor (Epoxomycin SIGMA 5μM, Syl-A 10μM, and MG-132 50μM SIGMA) incubated in a Percival growth cabinet for the desired time. Alternatively, inflorescences were dipped in a solution made of water, inhibitor at the desired concentration, 0.02% Tween-20. Both methods gave similar results.

### EdU staining

Inflorescences were cut leaving a 1.5cm stem from the apex, and developing siliques and fertilized flowers were removed with a vertical movement in order to peel away strips of epidermis. We observed that such scarification of the stem was necessary to ensure efficient EdU uptake. Inflorescences were then placed in a 1.5ml Eppendorf tube filled with MS liquid media (1/2 MS salt base [Carolina Biologicals, USA], 1% sucrose, 0.05%MES, pH>5.7 with KOH). We created a small hole in the tube lid where we inserted the inflorescence, so to avoid evaporation of the media. After that, we applied to the samples a gentle vacuum (2min with the vacuum valve slightly open). Tiny air bubbles should be visible on the submerged part of the stem. This step is fundamental to ensure efficient EdU uptake. The samples were then placed in a Percival growth cabinet and left overnight to recover. The morning after, the inflorescences were moved to a new 1.5ml tube filled with EdU-MS media (1/2 MS salt base [Carolina Biologicals, USA], 1% sucrose, 0.05%MES, pH>5.7 with KOH 200ml of MS 2% sucrose with 5mgr EdU), and incubated for the desired time in the Percival cabinet. Mature pistils were manually pollinated early morning for the time-course experiment. At the desired time, carpels were collected, sliced open along the replum, and incubated in fixative (Paraformaldehyde 4% in PBS1X pH7.4) for 30min at 4°C applying an initial 2 min vacuum, and then washed twice with PBS-Tween^0.1%^. Where required, samples were kept in PBS1X over/night at 4°C. Samples were then incubated in Digestion mix (cellulase 0.5%, pectolyase 0.5%, driselase 1%), for 1.5 hours at 37°C, and washed three times with PBS-Tween^0.1%^. Samples were then incubated for 1.5 hours 4°C in PBS1X-Tween^2%^, and then washed three times with water. The samples were then gently moved in a new clean tube filled with 100μl of Copper solution (Copper sulfate 6mM, 3μl of TEG-Azide, 2% Tween-20). 100μl of Sodium Ascorbate 30mM were squirted over by placing the Eppendorf tube on a vortex at low speed. Samples were then incubated in the completed Click-reaction for 1h at RT in the dark. After that, samples were rinsed several times with abundant water (at least 10 washes of 10 min each, until the samples do not leak yellow anymore), mounted in anti-fade vectashield, and imaged at a Leica SP5 following settings for AlexaFluor 488.

### Propidium Iodide staining

Propidium Iodide staining for ploidy quantification has been performed accordingly to She et al., 2014^38^. Briefly,x pistils were gently sliced along the replum and incubate in BVO buffer (2mM EGTA pH7.5, 1% formaldehyde, 10% DMSO, 0.1% Tween-20) for 30 min at RT on an orbital shaker. After that, samples were washed once with PBS1X-Tween^0.1%^, incubate 5 min in methanol, 5 min in ethanol, and stored at −20°C or processed immediately. Samples were then incubate in ethanol:xylene 1:1 mixture for 30min at RT, rinsed with ethanol for 5 min, and then with methanol for 5 min. Samples were then fixed in Fixative 2 (methanol:PBS-Tween^0.1%^ 1:1, 2.5% formaldehyde), for 15min at RT, and washed twice with PBS-Tween^0.1%^. Samples were then incubated in the Digestion mix (cellulase 0.5%, pectolyase 0.5%, driselase 1%), for 2 hours at 37°C. We recommend to optimize the digestion time for each new batch of digestion mix. After that, samples were washed twice with PBS-Tween^0.1%^, and incubated in PBS1X-Tween^1%^ supplemented with RNase (Qiagen) at a final concentration of 100ugr/ml for 1.5hours at 37’C. Samples were then washed twice with PBS1X-Tween^0.1%^, and fixed in Fixative 3 (2.5% formaldehyde, PBS1X-Tween^0.1%^), for 20min at RT, and washed twice with PBS-Tween^0.1%^. The samples were then incubated 2 hours at 4°C in PBS1X-Tween^2%^, washed twice with PBS-Tween^0.1%^, and incubated with Propidium iodide 10ugr/ml in PBS1X for 15 min at RT in the dark, washed again twice with PBS1X and mounted on a slide in Vectashield supplemented with Propidium iodide (Reactolab, Vector H-1300). Samples were imaged with a Leica TCS SP8 Multiphoton microscope, equipped with RLD detectors, and in photon counting mode.

### *In situ* hybridization

The *YFP* digoxygenin-labelled antisense and sense RNA probes were generated by *in vitro* transcription according to the instructions provided with the DIG RNA labelling kit (SP6/T7; Roche) using a plasmid containing the YFP sequence as template. Developing inflorescences or pollinated pistils and collected at the desired time points were fixed in FAA fixative (Formaldehyde, acetic acid, ethanol) o/n, then moved in ethanol 70% and into an ASP200 embedding machine for embedding (Leica Microsystems GmbH, Wetzlar, Germany), where they were dehydrated in a graded series of ethanol at room temperature (70% for 1 h, 3×90% for 1h, 3×99.98% for 1h, all at room temperature) followed by xylol (2×1 h and 1×1 h 15 min). Xylol was substituted by Paraplast X-tra embedding media (Roth AG, Arlesheim, Switzerland) at 58°C (2×1 h and 1×3 h). The pistils were poured into small plastic tray while the paraffin was still liquid, and then let to set. For storage, the paraffin blocks containing the pistils have been stored at 4°C. The samples were then sliced in 8-μm sections with a RM2145 Leica microtome (Leica Microsystems GmbH, Wetzlar, Germany) and then hybridized as described previously^72^, with strong formaldehyde washes^73^, as described in Dreni et al. 2007^74^. Images were taken with a Zeiss DMR microscope equipped with differential interference contrast (DIC) filters and Leica Flexacam C3 LSR camera.

### Clearing

Siliques at different time points after pollination were fixed o/n in fixative (Ethanol:Acetic Acid 9:1 v/v) at room temperature. The following day, the fixative was replaced with Ethanol 70%. Seeds and ovules were isolated from the valves and mounted in Clearing solution (Chloral Hydrate:Water:Glycerol 4:2,5:1 w/v/w) and left to clear few hours or overnight depending on the size. Images were taken with a Zeiss DMR microscope equipped with differential interference contrast (DIC) filters and Leica Flexacam C3 LSR camera.

### Pollen tube speed assay

Emasculated pistils were emasculated and pollinate the day after with pollen from the desired background. After 5 hours ovule were dissected and mounted on glucose 7% Propidium iodide 0.1mgr/ml. Ovule have been imaged at DM6000 Leica microscope by scoring the percentage of ovules with accumulation of Propidium Iodide in the synergid, a sign of occurred fertilization.

### Laser Capture Microdissection Transcriptome

The protocol is based on the method published in Schmidt et al., 2012^35^. Briefly, pistils obtained from the reciprocal crosses at precise time points were collected and fixed in ice-cold ethanol:acetic acid 9:1, and by applying vacuum for the initial 5 min by keeping the samples on ice. Afterwards, the samples were incubated in the fixative overnight at 4°C. For long storage of the material, fixative have been replaced by with ethanol 100%, and samples were stored at −20°C. For embedding, samples were transferred in ethanol 70%, and moved in an ASP200 embedding machine (Leica Microsystems GmbH, Wetzlar, Germany), where the tissue was dehydrated in a graded series of ethanol at room temperature (70% for 1 h, 3×90% for 1h, 3×99.98% for 1h, all at room temperature) followed by xylol (2×1 h and 1×1 h 15 min). Xylol was substituted by Paraplast X-tra embedding media (Roth AG, Arlesheim, Switzerland) at 58°C (2×1 h and 1×3 h). The pistils were poured into small plastic tray while the paraffin was still liquid, and then let to set. For storage, the paraffin blocks containing the pistils have been stored at 4°C.

For microdissection, paraffin blocks containing the pistils were cut on a RM2145 Leica microtome (Leica Microsystems GmbH, Wetzlar, Germany) to 8um thin slices and mounted on nuclease-free membrane-mounted metal-frame slides using DEPC water. Slides were left to dry for 1 hour on a heating block at 38-40°C. Samples were then deparaffinized in xylol at room temperature (2 washes of 20min each). Microdissection was performed using a mmi CellCut Plus device (MMI Molecular Machines & Industries AG, Glattbrugg, Switzerland). An average of 40 Central Cells were isolated per day using the special glued-cap Eppendorf tube. After collection, 50μl of extraction buffer of the PicoPure RNA isolation kit (Arcturus Engineering, Mountain View, USA) was added to the Eppendorf tube and incubated up-side down (so to have the buffer in contact with the cap of the tube) for 30min at 42°C in a water bath to release the tissue from the membrane. At intervals, the sample was vortexed vigorously by keeping the Eppendorf tube up-side down so to expose the sections to the buffer. The tube was then stored at – 80’C for longer storage. Total RNA was isolated using the PicoPure RNA isolation kit (Arcturus Engineering, Mountain View, USA) following the manufacturer’s instructions. Extracts from different tubes were pooled to reach a sufficient RNA yield. We have combined together tubes to obtain an average of 300 Central Cells per each replicate.

Total RNA was tested for integrity and quality using the Agilent TAPE Station, and converted to cDNA using the SMART-Seq^®^ v4 Ultra^®^ Low Input RNA Kit for Sequencing (Takara Bio), following the manufacturer’s instruction. Then libraries were constructed using the Nextera XT DNA Sample Preparation Kit (Illumina) following manufacturer’s instructions.

Libraries were then sequenced at 150bp double paired-end, in a Nova-Seq Illumina machine, by combining all the 12 libraries in a single lane at the Functional Genomic Centre Zurich.

For standard differential expression, short reads generated in this study were deposited at NCBI Sequence Read Archive (SRA, www.ncbi.nlm.nih.gov/sra) and are accessible through the accession number XXX. Reads were trimmed and quality-checked with fastp^75^ (version 0.20.1). Transcripts were quantified with Salmon^76^ (version 1.4.0) using the cDNA and gene annotation available from Araport (Araport 11). Variation in read counts was analysed with a general linear model in R (version 3.6.1) with the package DESeq2^77^ (version 1.24.0) according to a design with a single factor with multiple levels. Specific conditions were then compared with linear contrasts. Within each comparison, P-values were adjusted for multiple testing (Benjamini-Hochberg), and regions with an adjusted P-value (false discovery rate, FDR) below 0.01 and a minimal log2 fold-change (i.e., the difference between the log2 transformed, normalized sequence counts) of 2 was considered as differentially expressed. Normalized sequence counts were calculated accordingly with DESeq2 and log2(x+1) transformed.

To functionally characterize a gene set of interest, we tested for enrichment of gene ontology (GO) terms with topGO^78^ (version 2.32.0). Analysis was based on gene counts (genes in the set of interest compared to all annotated genes) using the “weight”“ algorithm with Fisher’s exact test (both implemented in topGO). A term was identified as significant if the P-value was below 0.05.

## Lead Contact

Further information and requests for resources and reagents may be directed to and will be fulfilled by Ueli Grossniklaus.

## Materials Availability

All new materials generated in this study will be available upon request from Ueli Grossniklaus.

## Data and Code Availability

The RNA-Seq raw data have been deposited at ArrayExpress under accession number XXX.

## FIGURE LEGENDS

**Extended Data Figure 1.**
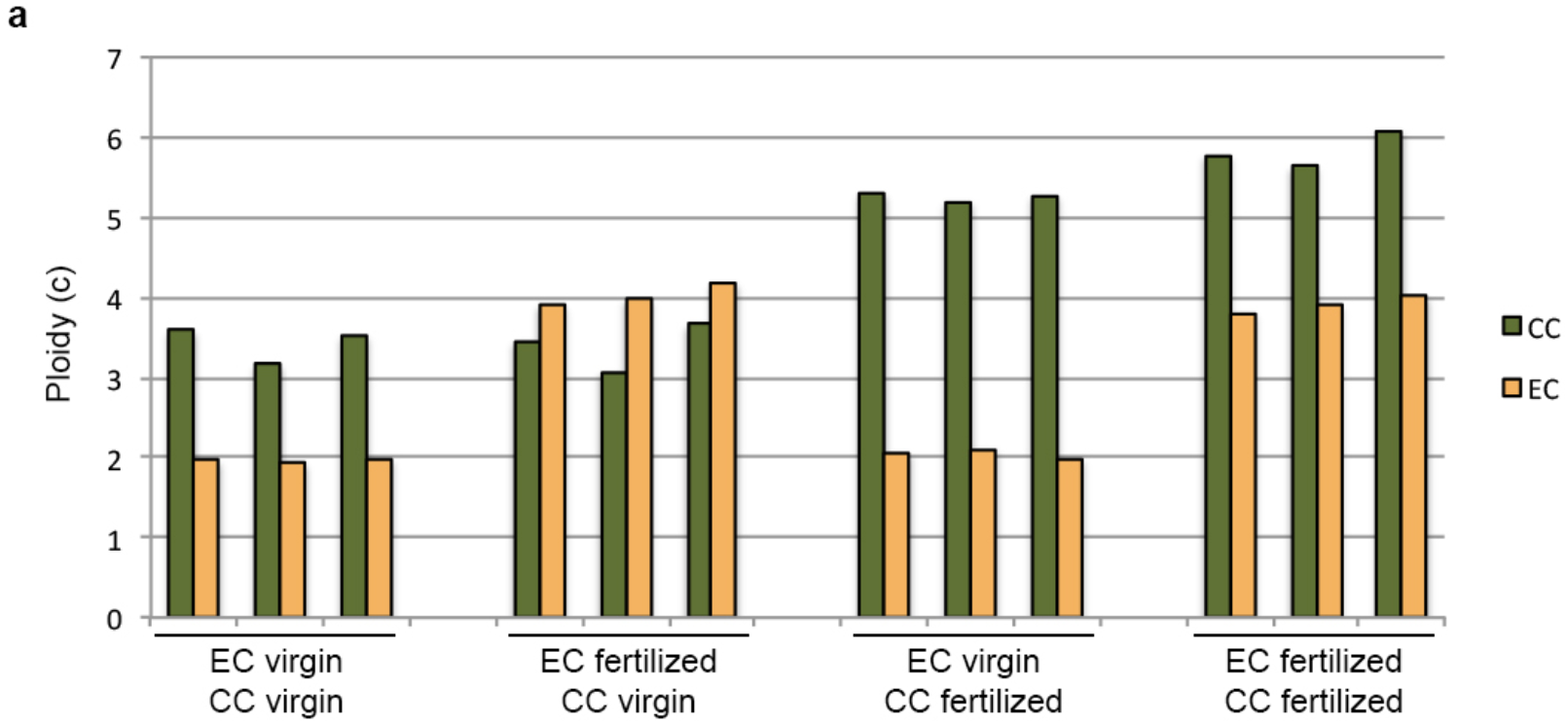
**a.** Ploidy of CC (green bars) and EC (yellow bars) at different time points.

**Extended Data Figure 2.**
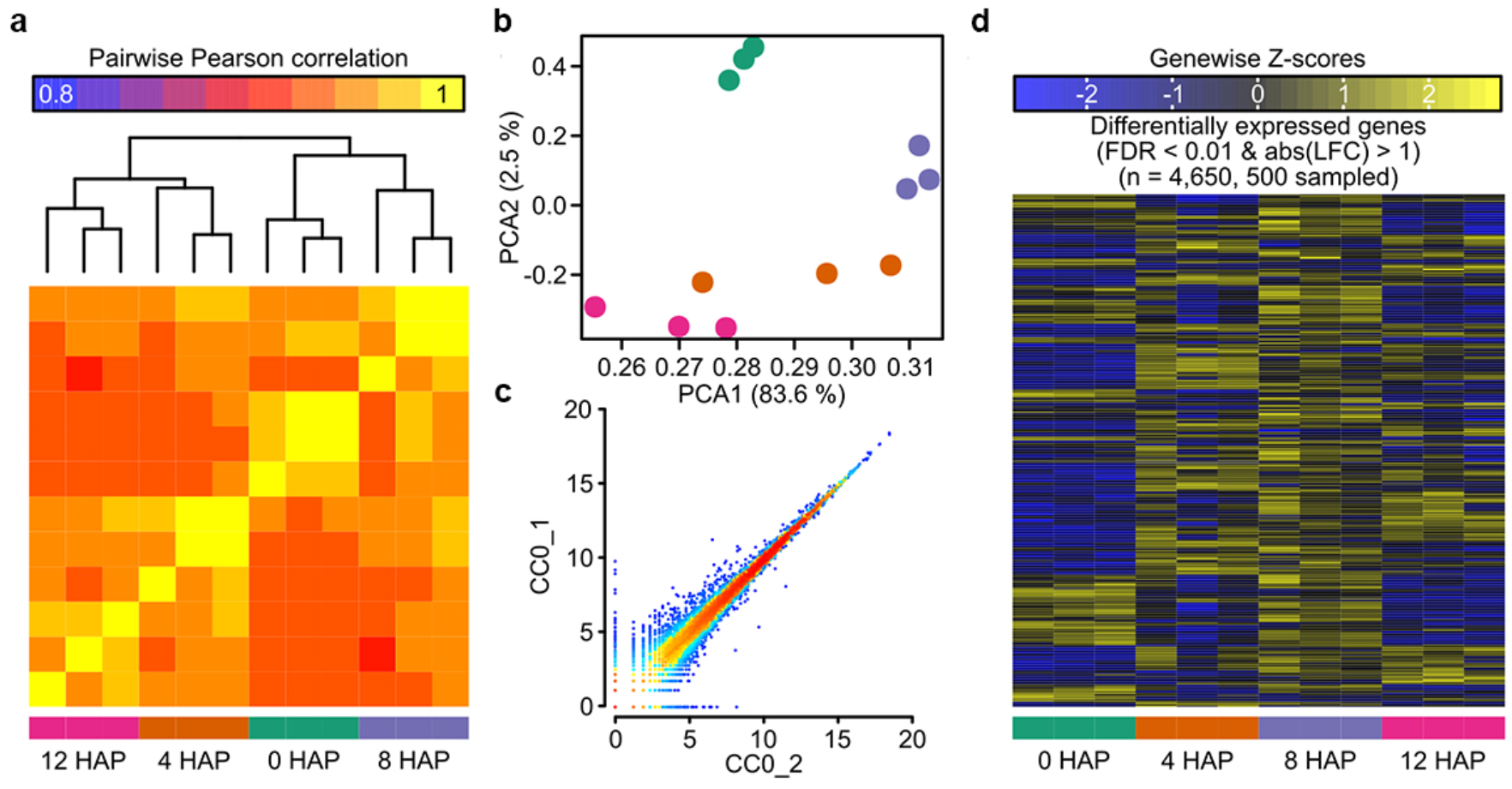
**a.** Pair-wise Pearson correlation coefficients and hierarchical clustering of samples after read count normalization. **b.** PCA of all samples using the normalized read counts. The first two components explain 86.1% of the variation in the data set. **c.** Representative pairwise scatter plot between two replicates. **d.** Heatmap with differentially expressed genes, a random subset of 500 genes is shown.

**Extended Data Figure 3.**
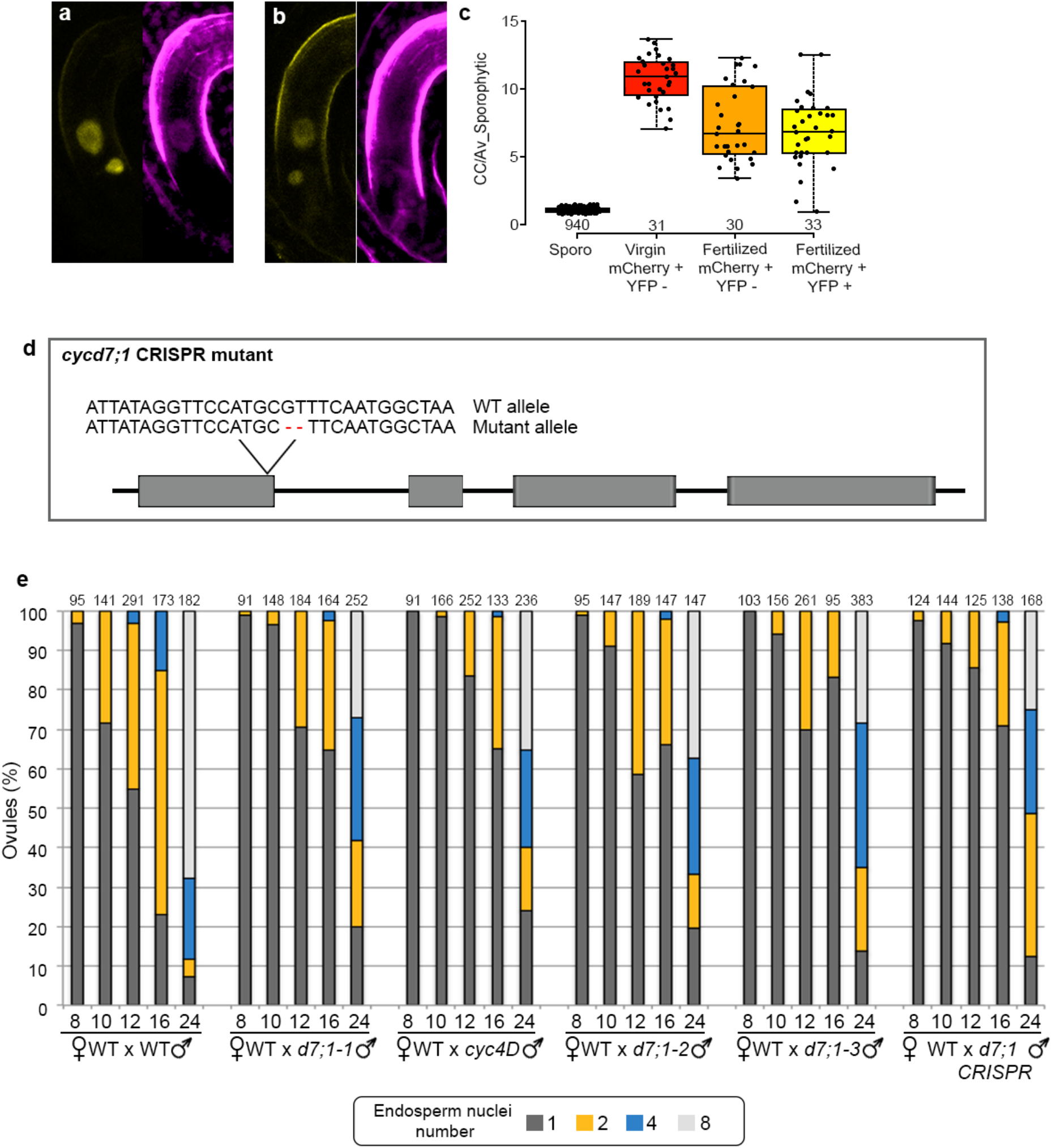
**a-b.** Confocal images of crosses between *pRBR1::mCherry-RBR1* with *pCYCD7;1::CYCD7;1-YFP* showing presence of RBR1-mCherry and CYCD7;1-YFP in the CC (a), and absence of RBR1-mCherry with CYCD7;1-YFP presence (b). **c.** Box plot showing the quantification of *pRBR1::mCherry-RBR1* with *pCYCD7;1::CYCD7;1-YFP* signal in fertilized ovules. **d**. Schematic representation of the CRISPR/Cas9 allele created in *CYCD7;1*. **e**. Classification of fertilized ovules (in percentage) based on the number of endosperm nuclei at a certain time point (8, 10, 12, 16, 24 HAP), in crosses between *WT x WT, WT x cycd7;1-1, WT x cyc4D, WT x cycd7;1-2, WT x cycd7;1-3*, and *WT x cycd7;1-CRISPR*. Sample size is indicated at the top of each bar.

**Extended Data Figure 4.**
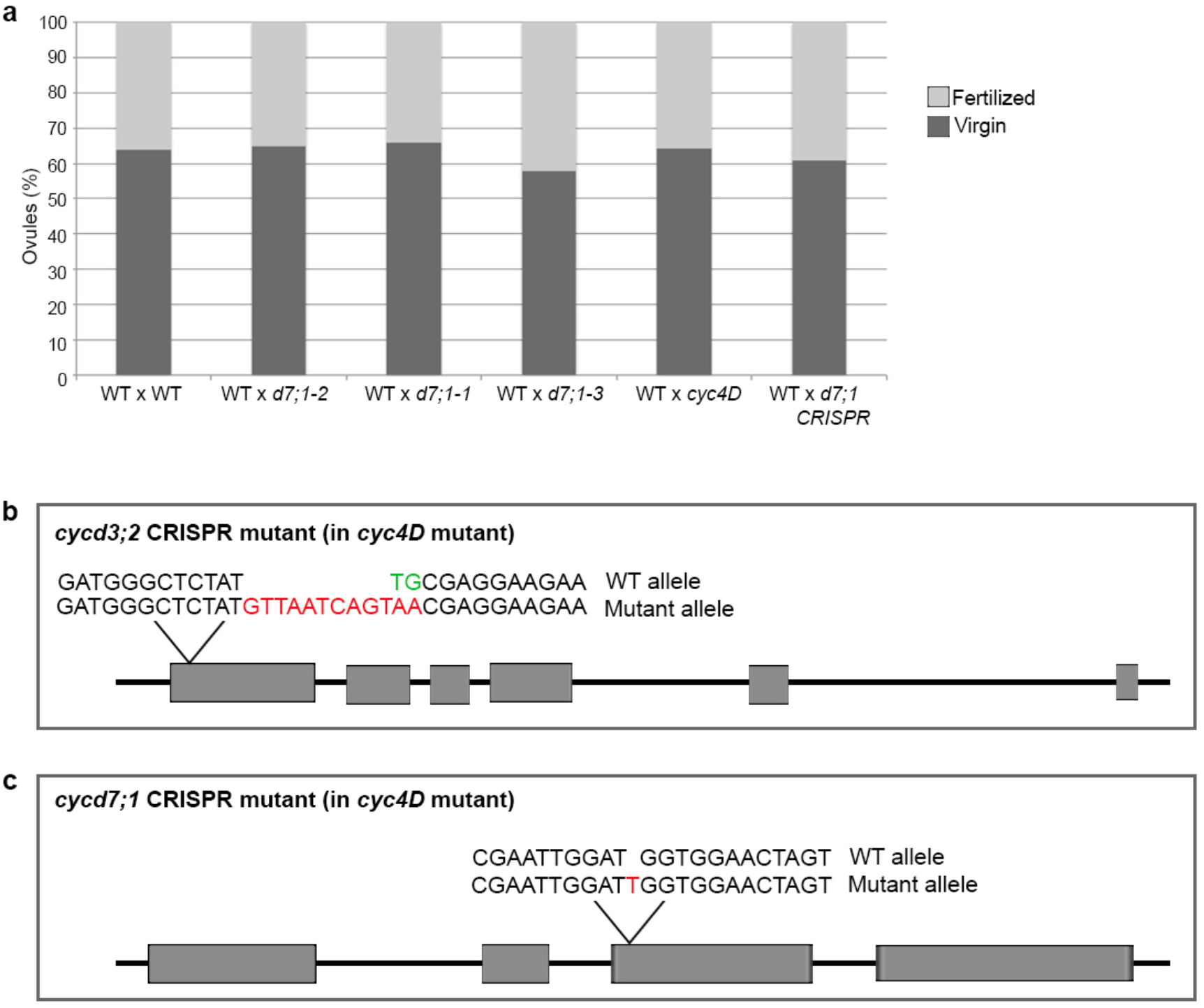
**a.** Chart of virgin and fertilized ovule to assess pollen tube growth of various mutant pollen compared to the wild type. **b-c**. Schematic representation of the CRISPR/Cas9 allele created in *CYCD7;1* and *CYCD3;2* to create the quadruple *cycd3;1 cycd3;2 cycd3;3 cycd7;1* mutant (*cyc4D*).

